# Monosynaptic inputs to ventral tegmental area glutamate and GABA co-transmitting neurons

**DOI:** 10.1101/2023.04.06.535959

**Authors:** Emily D. Prévost, Alysabeth Phillips, Kristoffer Lauridsen, Gunnar Enserro, Bodhi Rubinstein, Daniel Alas, Dillon J. McGovern, Annie Ly, Makaila Banks, Connor McNulty, Yoon Seok Kim, Lief E. Fenno, Charu Ramakrishnan, Karl Deisseroth, David H. Root

## Abstract

A unique population of ventral tegmental area (VTA) neurons co-transmits glutamate and GABA as well as functionally signals rewarding and aversive outcomes. However, the circuit inputs to VTA VGluT2+VGaT+ neurons are unknown, limiting our understanding of the functional capabilities of these neurons. To identify the inputs to VTA VGluT2+VGaT+ neurons, we coupled monosynaptic rabies tracing with intersectional genetic targeting of VTA VGluT2+VGaT+ neurons in mice. We found that VTA VGluT2+VGaT+ neurons received diverse brain-wide inputs. The largest numbers of monosynaptic inputs to VTA VGluT2+VGaT+ neurons were from superior colliculus, lateral hypothalamus, midbrain reticular nucleus, and periaqueductal gray, whereas the densest inputs relative to brain region volume were from dorsal raphe nucleus, lateral habenula, and ventral tegmental area. Based on these and prior data, we hypothesized that lateral hypothalamus and superior colliculus inputs were glutamatergic neurons. Optical activation of glutamatergic lateral hypothalamus neurons robustly activated VTA VGluT2+VGaT+ neurons regardless of stimulation frequency and resulted in flee-like ambulatory behavior. In contrast, optical activation of glutamatergic superior colliculus neurons activated VTA VGluT2+VGaT+ neurons for a brief period of time at high stimulation frequency and resulted in head rotation and arrested ambulatory behavior (freezing). For both pathways, behaviors induced by stimulation were uncorrelated with VTA VGluT2+VGaT+ neuron activity. However, stimulation of glutamatergic lateral hypothalamus neurons, but not glutamatergic superior colliculus neurons, was associated with VTA VGluT2+VGaT+ footshock-induced activity. We interpret these results such that inputs to VTA VGluT2+VGaT+ neurons may integrate diverse signals related to the detection and processing of motivationally-salient outcomes. Further, VTA VGluT2+VGaT+ neurons may signal threat-related outcomes, possibly via input from lateral hypothalamus glutamate neurons, but not threat-induced behavioral kinematics.

## INTRODUCTION

Dopamine-, GABA-, and glutamate-releasing ventral tegmental area (VTA) neurons regulate rewarding and aversive experiences as well as the anticipation of these outcomes by learned predictors (Schultz, 1998; Lammel et al., 2011; Cohen et al., 2012; Tan et al., 2012; van Zessen et al., 2012; Ilango et al., 2014; Root et al., 2014a; Wang et al., 2015; Qi et al., 2016; Yoo et al., 2016; Morales and Margolis, 2017; Root et al., 2018a). From their initial discovery (Yamaguchi et al., 2007), VTA glutamate-releasing neurons were highlighted for their molecular heterogeneity that reflects unique capabilities to release different combinations of neurotransmitters. In particular, subsets of VTA glutamate neurons release glutamate alone (requiring the vesicular glutamate transporter 2, VGluT2), glutamate with dopamine (requiring VGluT2 and tyrosine hydroxylase, TH), or glutamate with GABA (requiring VGluT2 and the vesicular GABA transporter, VGaT) (Yamaguchi et al., 2007; Stuber et al., 2010; Yamaguchi et al., 2011; Tritsch et al., 2012; Root et al., 2014b; Zhang et al., 2015; Root et al., 2018b; Root et al., 2020). In contrast to other VTA cell-types, VTA VGluT2+VGaT+ neurons that release both glutamate and GABA show a unique profile of neuronal activity consisting of activation by rewarding or aversive outcomes but not by learned predictors of those outcomes (Root et al., 2020).

The circuit inputs that regulate VTA VGluT2+VGaT+ neuronal signaling are unknown, limiting our understanding of how their unique outcome-signaling properties arise. To begin identifying the circuit inputs to VTA glutamate and GABA cotransmitting neurons, we first performed whole-brain monosynaptic rabies tracing of retrogradely-labeled neurons from VTA VGluT2+VGaT+ neurons and utilized SHARCQ to register traced neurons to the Allen Brain Atlas (Lauridsen et al., 2022). Brainwide inputs largely arose from regions linked with threat, action selection, outcome valuation, and motor-related signaling. Within threat-related circuits, we identified that glutamatergic projections from the lateral hypothalamus (LH) and superior colliculus (SC) increased VTA VGluT2+VGaT+ neuronal activity *in vivo*. Based on our observations during stimulation of these neurons, we performed an initial exploration of stimulation-induced behaviors. Activation of glutamatergic LH and SC neurons resulted in distinct threat-related behavioral repertoires that were not correlated with VTA VGluT2+VGaT+ neuron activation, suggesting VGluT2+VGaT+ neurons are capable of signaling a generalized threat regardless of the motor responses that underlie them. However, glutamatergic lateral hypothalamus activation of VTA VGluT2+VGaT+ neurons was selectively associated with VGluT2+VGaT+ footshock-related activity, suggesting this input provides threat-related information to these neurons. Together, these results provide novel insights into the cell-type specific influences on VTA VGluT2+VGaT+ neuronal activity in the integration of motivationally salient outcome-signaling.

## METHODS

### Animals

VGluT2-IRES::Cre mice *(Slc17a6*^*tm2(cre)Lowl*^/J; Jax Stock #016963) were crossed with VGaT-2A::FlpO mice (*Slc32a1*^*tm2*.*1(flpo)Hze*^/J; Jax Stock #031331) at the University of Colorado Boulder to produce VGluT2::Cre/VGaT::FlpO offspring. Mice were maintained in a colony room with a 12 h light/dark cycle (lights on at 7:00 h) with *ad libitum* access to food and water. All animal procedures were performed in accordance with the National Institutes of Health Guide for the Care and Use of Laboratory Animals and approved by the University of Colorado Boulder Institutional Animal Care and Use Committee.

### Stereotactic surgery

Male and female VGluT2::Cre/VGaT::FlpO mice (*N* = 8, 12-20 weeks of age) were anesthetized with 1-3% isoflurane and secured in a stereotactic frame (Kopf). INTRSECT vectors encoding optimized rabies glycoprotein (oG) and an avian cell-surface receptor tethered to mCherry (TVA-mCherry) were used for targeting VGluT2+VGaT+ neurons (Fenno et al., 2014; Hafner et al., 2019; Fenno, 2020). AAV8-nEF-Con/ Fon-TVA-mCherry (Addgene 131779, 5×10^12^ titer) and AAV8-EF1α-Con/Fon-oG (Addgene 131778, 5×10^12^ titer) were mixed in equal proportions (final titer = 2.5×10^12^ each) and injected into VTA (AP: −3.2 mm relative to bregma; ML: 0.0 mm relative to midline; DV: −4.4 mm from skull surface; 500 nL). Three weeks later, mice were injected at the same coordinates with SAD-ΔG-EGFP(EnvA) (Salk, 400 nL). Total injection volume and flow rate (100 nL/min) were controlled with a microinjection pump (Micro4; World Precision Instruments, Sarasota, FL). The syringe was left in place for an additional 10 min following injection to allow for virus diffusion, after which the syringe was slowly withdrawn to prevent spread by capillary action. Brains were harvested 1 week after rabies injection as described in *Histology*. Control mice (*N* = 4 VGluT2::Cre mice and *N* = 4 VGaT::Flp mice) were injected in VTA with the same Cre and Flp-dependent TVA and oG AAVs, followed three weeks later by injection of SAD-ΔG-EGFP(EnvA) in the same site. Surgical, histological, and analytical procedures were identical to VGluT2::Cre/VGaT::FlpO mice.

For *in vivo* optical stimulation and photometry experiments, male and female VGluT2::Cre/ VGaT::FlpO mice were injected with AAV8-EF1-Con/Fon-GCaMP6m (Addgene 137119, 8.5×10^12^ titer, 500 nL) into VTA (AP: −3.2; ML: 0.0; DV: −4.4 mm; 500 nL). In the same surgery, AAV8-nEF-Con/Foff-ChRmine-oScarlet (Addgene 137161, 1×10^13^ titer) was bilaterally injected into either LH (AP: −1.2; ML: ±1.2; DV: −5.0; 400 nL; *N* = 4 mice) or SC (AP: −3.5; ML: ±0.8; DV: −1.9 mm; 500 nL; *N* = 4 mice). All mice were implanted with a recording fiber cannula (400 μm core diameter, 0.66 NA; Doric Lenses, Québec, Canada) dorsal to VTA (AP: −3.2 mm relative to bregma; ML: +1.0 mm at 9.5° relative to midline; DV: −4.2 mm from skull surface). Additionally, mice were implanted with optic fiber cannulae (200 μm core diameter, 0.37 NA; Doric Lenses) in either LH bilaterally (AP: −1.2; ML: −2.0 at 10° [left] and +2.9 at 20° [right]; DV: −4.7 [left] and −4.9 [right]) or superior colliculus unilaterally (AP: −3.5; ML: −1.1 at 10°; DV: −1.73 mm). The LH stimulating implants in left and right hemisphere were angled differently to better avoid the recording VTA implant on the skull. All implants were secured to the skull with screws and dental cement. Mice were allowed to recover for three weeks before experimentation.

### Pavlovian fear conditioning

Wildtype C57Bl6 mice and GCaMP mice were brought over four days to behavior chambers outfitted with rod flooring which was electrically connected to a shock generator (Med-Associates). Mice were exposed to 5 auditory conditioned stimuli per day (CS+: 10-kHz tone, 30 s) that co-terminated with the delivery of footshock (US: 0.5 s, 0.5mA, onset 29.5 s after cue onset). Cues were presented on a variable interval 60-second schedule. On the fourth day, GCaMP6m signal was recorded. For wildtype mice, AnyMaze tracked the location of mice (30 Hz) and quantified freezing 5 seconds before and during each conditioned stimulus. Freezing parameters in AnyMaze required at least 1 s of freezing to identify a freezing bout.

### ChRmine stimulation

Three weeks after surgery, mice were restricted by the tail with gauze tape and placed in an enclosed Plexiglas box (Med-Associates, MED-TLH-M). For frequency analysis, VTA GCaMP6m signals were recorded while optically stimulating ChRmine (589 nm, 10 ms pulse duration, 10-15 mW) at 5, 10, 20, and 40 Hz (Marshel et al., 2019). Five trials were delivered at each frequency. Each laser train was one second in duration, delivered on a 60-second inter-train interval and five minutes between frequencies. Frequency order was randomly assigned. After at least three days, mice were tail-restricted with gauze tape in an unenclosed space allowing for rotational, ambulatory, or rearing movement that was behaviorally analyzed. ChRmine was activated at the maximum GCaMP frequency response for each input (20 Hz in LH and 40 Hz in SC, 10-15 mW) using 589 nm light at 10 ms pulse duration and one-second train duration. Ten trains were delivered with one-minute inter-train intervals. Video recordings were made concurrently with ChRmine stimulation (TDT, 30 Hz).

### GCaMP recordings

GCaMP6m was excited at 465 nm and 405 nm (isosbestic control) with amplitude-modulated signals from two light-emitting diodes reflected off dichroic mirrors and coupled into an optic fiber. Signals from GCaMP and the isosbestic control channel were returned through the same optic fiber and acquired with a femtowatt photo-receiver (Newport, Irvine, CA), digitized at 1 kHz, and recorded by a real-time signal processor (Tucker Davis Technologies, TDT). Analysis of the recorded calcium signal was performed using custom-written MATLAB scripts. Signals (465 nm and 405 nm) were downsampled (10x) and peri-event time histograms (PETHs) were created trial-by-trial between −10 sec to +30 sec surrounding each optical stimulation train onset and −10 sec to +10 sec surrounding conditioned stimulus (CS+) or footshock onsets. For each trial, data was detrended by regressing the isosbestic control signal (405 nm) on the GCaMP signal (465 nm) and then generating a predicted 405 nm signal using the linear model generated during the regression. The predicted 405 nm channel was subtracted from the 465 nm signal to remove movement, photo-bleaching, and fiber bending artifacts (dF). Baseline normalized maximum z-scores (normalized dF) were taken from −6 to −3 seconds prior to LH train, CS+, or footshock onset and maximum event z-scores were taken from 0 to 2 seconds following LH train, SC train, CS+, or footshock onset (McGovern et al., 2021; McGovern et al., 2023). Due to the shorter activation profile following superior colliculus stimulation, PETHs were created trial-by-trial between −5 sec and +5 sec surrounding each train onset and baseline normalized maximum z-score was taken from −3 to 0 seconds prior to superior colliculus train onset. Area under the curve (AUC) was collected between 0 and 2 sec following stimulation train onset.

### Video scoring

Behavioral optical stimulation experiments were video recorded at 30 Hz (TDT). For simultaneous optical stimulation and photometry experiments, foot treading, rearing, freezing, and turning behavior was timestamped in OpenScope (TDT). Foot treading was defined as more than one full gait cycle with all four paws. Rearing was defined as both forepaws lifting from the ground. Freezing was marked as the complete cessation of movement except that which is necessary for breathing. For superior colliculus VGluT2 ChRmine experiments, head turning was quantified by subtracting the absolute angle of the head at freeze onset from the absolute head angle immediately before stimulation onset. Angles were determined by placing a protractor parallel to the base of the apparatus with the origin positioned at the midpoint between the animal’s scapulae and extending a straight line to the bridge of the nose. Body turn calculations were performed the same way but with the origin of the protractor at the midpoint between the animal’s hindlegs and with the line extending to the midpoint of the scapulae. All angles are presented in absolute values. Supplemental movies were created with Canva.

### Histology

Mice were anesthetized with isoflurane and perfused transcardially with 0.1 M phosphate buffer followed by 4% (w/v) paraformaldehyde in 0.1 M phosphate buffer, pH 7.3. Brains were extracted, post-fixed for two hours in the same fixative, and cryoprotected in 18% sucrose in phosphate buffer at 4°C. For the tracing experiment, coronal sections of the whole brain (30 μm) were taken on a cryostat, mounted to gelatin-coated slides, and coverslipped with DAPI-containing Prolong Diamond medium. Sections were imaged on a PerkinElmer Opera Phenix High Content Screening System at 20X air objective magnification. Fluorescence images in the DAPI, EGFP, and mCherry channels were taken in a single plane, in which somata were in focus. For stimulation and photometry experiments, coronal sections containing VTA and either LH or SC (30 μm) were taken, mounted to gelatin-coated slides, and imaged for oScarlet expression, GCaMP6m expression, and optical fiber placement on a Zeiss Axioscope.

### Atlas registration and neuron quantification

Somata of EGFP-positive neurons were manually counted in FIJI using the Multi-point tool. Files of counted neurons’ X,Y positions within the image were exported using the Measure function. Somata of EGFP-mCherry co-expressing neurons were manually counted in Photoshop using the Count tool. Files of counted neurons’ X,Y positions were exported using the custom-written get_xy_photoshop.js JavaScript script (available at https://www.root-lab.org). Histological images and files of counted neurons’ locations were registered to the Allen mouse brain atlas using the open-source software tool SHARCQ (https://github.com/wildrootlab/SHARCQ) (Lauridsen et al., 2022). To avoid oversampling, coronal sections within 0.1 mm anteroposterior position of each other were compared, and the section with lower tissue integrity or imaging quality was removed from analysis. SHARCQ analysis with the Allen Atlas CCFv3 (Wang et al., 2020) localized 13.2% (8,756) of input neurons and 26.7% (606) of starter neurons simply to hypothalamus, midbrain, pallidum, or pons with no further subregion specification. The registered coordinates of those neurons were analyzed in the Kim Atlas^28^ to further classify neurons into more specific regions. Including both Allen Atlas and Kim Atlas data, neurons were categorized into 1,387 distinct regions which were merged into 556 regions in post-SHARCQ analysis.

Based on manual inspection, starter neurons designated to cerebellar peduncles, crossed tectospinal pathway, dorsal tegmental decussation, Edinger-Westphal nucleus, epithalamus related, extrapyramidal fiber systems, hypothalamic lateral zone, hypothalamus related, interpeduncular fossa, mammillary peduncle, mammillotegmental tract, medial forebrain bundle, medial lemniscus, midbrain, midbrain motor related, midbrain raphe nuclei, midbrain reticular nucleus, nucleus of Darkschewitsch, oculomotor nerve, oculomotor nucleus, periaqueductal gray, posterior hypothalamic nucleus, red nucleus, rubrospinal tract, substantia nigra compact part, tectospinal pathway, and ventral tegmental decussation were examined by hand and determined to belong to VTA. Likewise, starter neurons designated to lateral mammillary nucleus, mammillary body, mammillary related, and medial mammillary nucleus were determined to belong to the supramammillary nucleus. Input neurons localized to white matter tracts which are completely encapsulated by the VTA (dorsal tegmental decussation, mammillary peduncle, mammillotegmental tract, and ventral tegmental decussation) were moved post-hoc to the VTA category. Neurons localized to the superior cerebellar peduncles posterior to −4.80 mm relative to bregma were moved post-hoc to the parabrachial nucleus.

For density analysis, brain region volumes were calculated by summing the voxels in each region and converting to mm^3^ according to the voxel size of the atlas annotation (10 µm). The density of input neurons per brain region was calculated in neurons/mm^3^. Regions with fewer than 10 neurons on average were excluded from further density analysis.

### 3D rendering

The open-source Python package brainrender (Claudi et al., 2021) was used to generate 3D whole-brain renders and coronal slices of the Allen mouse brain atlas with neuron locations acquired in SHARCQ. Anatomical coordinates of starter or input neurons were combined across 8 animals for all plots. In 3D coronal slices, select brain structures were outlined for anatomical reference. Density heatmaps of starter and input regions were generated using the Matplotlib Python library and Seaborn KDE analysis according to the process described by the Wang lab (Takatoh et al., 2021). Anatomical coordinates of starter or input neurons in brain regions of interest were isolated. For bilateral structures, neurons of each hemisphere were plotted separately to prevent proximity algorithms from misrepresenting density across the midline. Density plots were overlaid onto 3D coronal atlas slice renders generated in brainrender. Plots were aligned in Adobe Photoshop using axes as guides, which were removed from the final image. Automatically generated scale bars indicate the color-coded density in neurons/mm^3^. Bilateral structures have discrete scale bars for each hemisphere. Parameters, code, and documentation are available at www.root-lab.org.

### Statistical testing

Statistical testing was performed in SPSS or GraphPad Prism. For LH VGluT2 or SC VGluT2 ChRmine stimulation experiments, a within-subjects ANOVA compared stimulation epoch (baseline, stimulation) and frequency (5, 10, 20, 40 Hz). Main effects were followed up by Sidak adjusted pairwise comparisons. A paired t-test compared the duration of reaction or freezing during baseline and stimulation. Simple linear regressions were used to correlate within-subject GCaMP activity (maximum Z-score or area under the curve from 0 to 2 sec) and parameters of the behavioral responses (reaction/freeze duration, latency to head turn, head turn angle, or latency to freeze). For superior colliculus VGluT2 ChRmine stimulation experiments, a paired t-test compared the average proportion of trials resulting in turn-and-freeze behavior in stimulation epochs versus inter-stimulation intervals. Within subjects, the normality of head turn and body turn angle distributions were tested using a Shapiro-Wilk test. A non-parametric Kruskal-Wallis test compared the latencies of head turn, peak GCaMP value, and freezing. The main effect was followed up by Dunn’s multiple comparison test. To examine CS+ and footshock-related activity, a within-subjects ANOVA compared the maximum baseline z-score (−6 to −3 sec from the CS+) with the maximum z-score following CS+ onset (0 to 2 sec) and the maximum z-score following footshock onset (0 to 2 sec). A main effect was followed up by contrast tests comparing baseline with CS+ and baseline with footshock (McGovern et al., 2021). For cue-induced freezing behavior in wildtype mice, a paired t-test compared percent of freezing 5 seconds before each cue and the initial 5 seconds of each cue. Pearson’s correlation was used to correlate VTA VGluT2+VGaT+ maximum z-score (0 to 2 sec) following 40 Hz stimulation of each pathway with footshock-induced maximum z-score (0 to 2 sec). For all tests, alpha was set to 0.05.

## RESULTS

We aimed to identify circuits that contribute to VTA VGluT2+VGaT+ neuronal signaling. To accomplish this, VGluT2::Cre/VGaT::Flp mice were injected in VTA with a mix of rabies helper viruses dependent on the expression of both Cre and Flp which encode a cell-surface avian receptor tethered to mCherry (TVA-mCherry) and optimized rabies glycoprotein (oG) (Watabe-Uchida et al., 2012; Kim et al., 2016; Hafner et al., 2019). Three weeks later, mice were injected in VTA with an avian-enveloped, glycoprotein-deleted rabies virus encoding EGFP (**Figure 1A**). After one additional week, brains were extracted, sectioned, imaged, counted, and registered to the Allen Atlas by SHARCQ (Lauridsen et al., 2022) (**Figure 1B**). 2,273 putative starter neurons were identified by coexpression of TVA-mCherry and rabies-EGFP (284.12 ± 47.51 neurons/mouse). The vast majority of putative starter neurons were found within the VTA (85.96 ± 2.50%) and a smaller subset were found within the supramammillary nucleus (14.04 ± 2.50%), consistent with the localization of midbrain VGluT2+VGaT+ neurons (Root et al., 2018) (**Figure 1C-J**). VTA starter neurons were highly dense within the interfascicular nucleus but also showed high density in the rostral linear nucleus and other portions of the VTA (**Figure 1H**).

**Figure 1.**
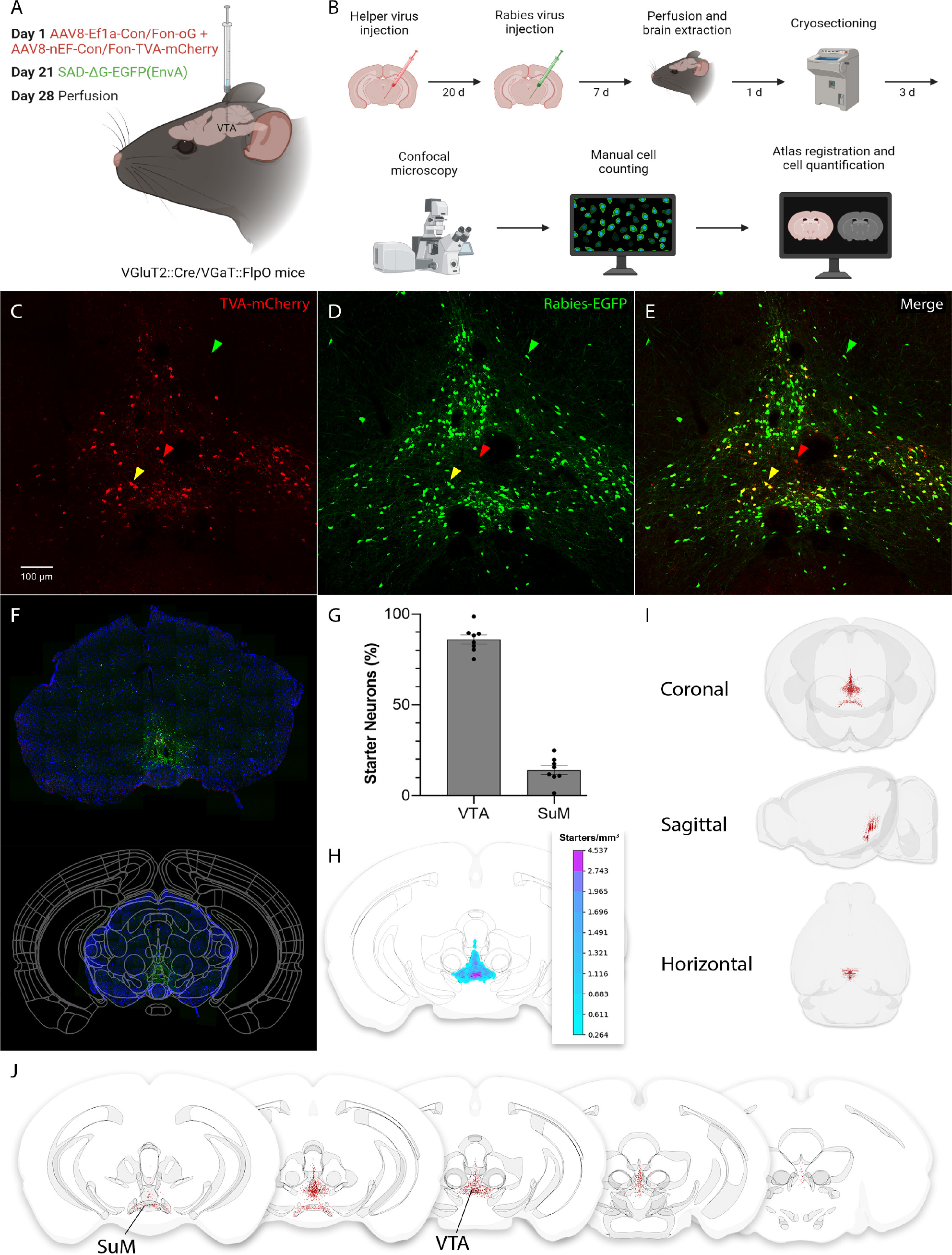
Experimental design and VTA VGluT2+VGaT+ starter neurons. **A**. Schematic of viral injections for retrograde tracing in VGluT2::Cre/VGaT::Flp mice. **B**. Timeline of viral injections, tissue processing, and analysis. **C-E**. Example fluorescence images of viral injections in the VTA. **C**. VGluT2+VGaT+ neurons expressing the helper viruses. **D**. Neurons expressing the modified rabies virus. **E**. Neurons with mCherry-EGFP overlap (yellow triangle) co-expressed the helper viruses and rabies virus, and comprised the population of starter neurons. Neurons with only mCherry expression (red triangle) did not receive the rabies virus, and thus were not included in the starter neuron population. Neurons with EGFP only (green triangle) expressed the rabies virus and comprised the population of monosynaptic input neurons. **F**. Example histological section (top) registered to the Allen mouse brain atlas using SHARCQ (bottom). **G**. Average proportion of starter neurons localized to the VTA and SuM. Dots represent individual animals, *N* = 8. **H**. Density heatmap of all starter neurons from −2.97 to −3.27 mm relative to bregma (*N* = 8 animals). **I**. Coronal, sagittal, and horizontal 3D representations of all starter neurons (*N* = 8 animals). **J**. 3D coronal slice representations of starter neurons from (left to right) −2.37 to −2.67, −2.67 to −2.97, −2.97 to −3.27, −3.27 to −3.57, and −3.57 to −3.87 mm relative to bregma (*N* = 8 animals). Abbreviations: SuM = supramammillary nucleus, VTA = ventral tegmental area.

To identify the intersectional specificity of the starter neurons, mice expressing Cre under the control of only the VGluT2 promoter (VGluT2::Cre) or mice expressing Flp under the control of only the VGaT promoter (VGaT::Flp) were first injected in VTA with Cre- and Flp-dependent AAVs encoding TVA-mCherry and oG. After three weeks, mice were injected in VTA with rabies-EGFP, and following an additional week examined for brainwide TVA-mCherry and rabies-EGFP expression (**Figure 2A**). In VGluT2::Cre mice, 11.25 ± 3.61 VTA neurons expressed rabies-EGFP, and in VGaT::Flp mice 3.25 ± 1.31 VTA neurons expressed rabies-EGFP (**Figure 2B**). In both cases we observed no mCherry expression nor were retrogradely-labeled rabies-EGFP neurons identified outside the injection site (**Figure 2C-D**). We interpret these data to indicate that rabies-EGFP neurons outside the VTA in VGluT2::Cre/ VGaT::Flp mice were retrogradely labeled from Cre and Flp co-expressing VTA neurons. However, a small number of rabies-EGFP neurons locally within the VTA may result from recombinase-independent labeling.

**Figure 2.**
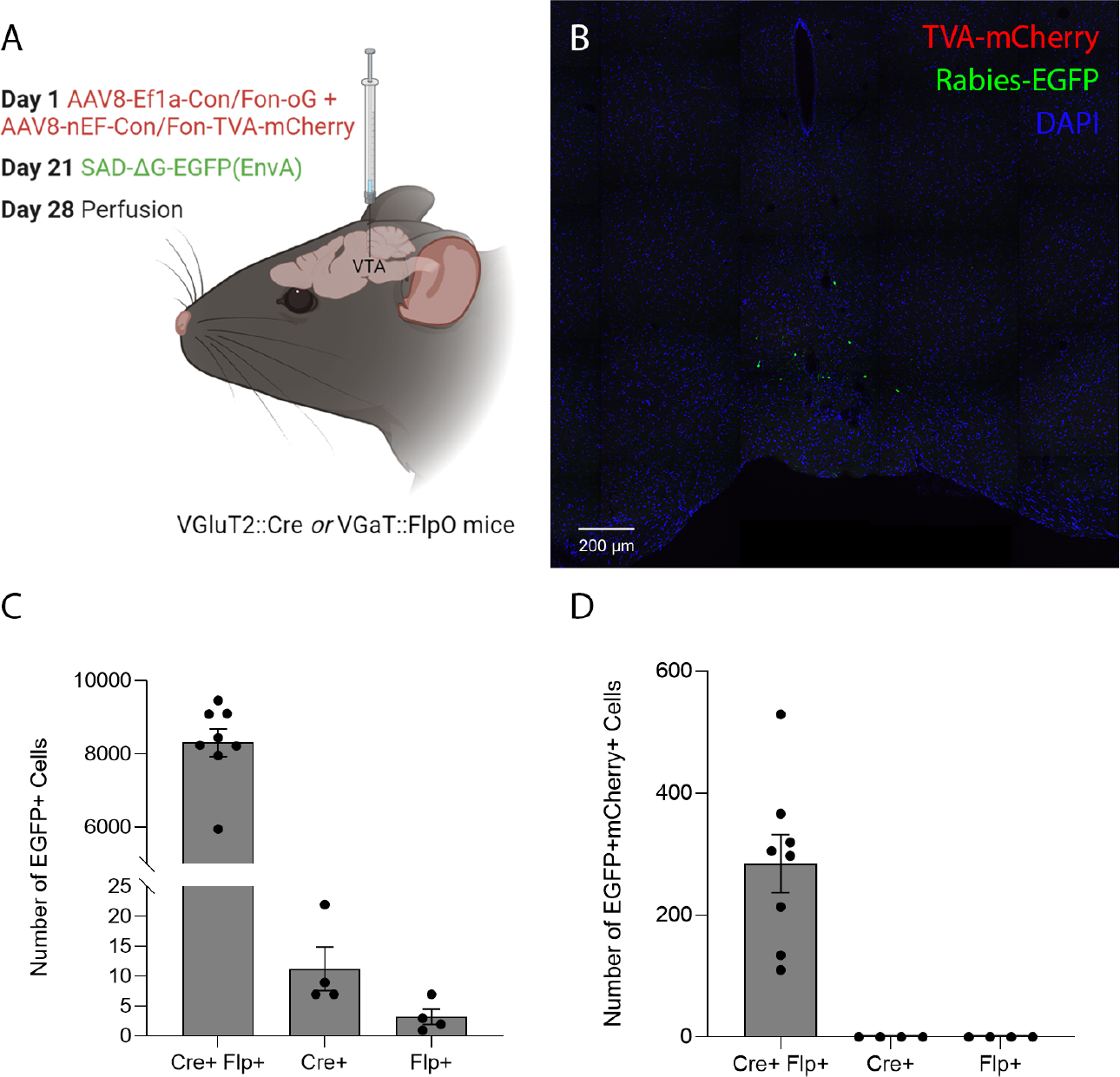
Intersectional control experiment. **A**. Schematic of viral injections for retrograde tracing in VGluT2::Cre or VGaT::Flp mice. **B**. Example fluorescence image of viral injection in the VTA. **C**. Number of EGFP-expressing VTA neurons in VGluT2::Cre/VGaT::Flp, VGluT2::Cre, or VGaT::Flp mice. **D**. Number of EGFP and mCherry co-expressing VTA neurons in VGluT2::Cre/VGaT::Flp, VGluT2::Cre, or VGaT::Flp mice.

Using monosynaptic rabies viral tracing of inputs to VGluT2+VGaT+ neurons, 66,465 input neurons (defined by EGFP-expression without mCherry-co-expression) were found to synapse onto VGluT2+VGaT+ neurons (8,308 ± 384.52 inputs per mouse). SHARCQ registration of these inputs to the Allen Atlas revealed high percentages of inputs in superior colliculus (7.69 ± 0.53%), midbrain reticular nucleus (6.24 ± 0.42%), periaqueductal gray (6.14 ± 0.32%), locally within the VTA (5.32 ± 0.29%), and LH (4.03 ± 0.37%) (**Figure 3A-C**). Considering input number relative to region volume, input neurons were densest in the dorsal raphe nucleus (1232.08 ± 161.26 neurons/mm^3^), lateral habenula (980.51 ± 114.79 neurons/mm^3^), locally within the VTA (713.74 ± 50.99 neurons/mm^3^), and supraoculomotor periaqueductal gray (480.86 ± 96.85 neurons/mm^3^) (**Figure 3D-E**).

**Figure 3.**
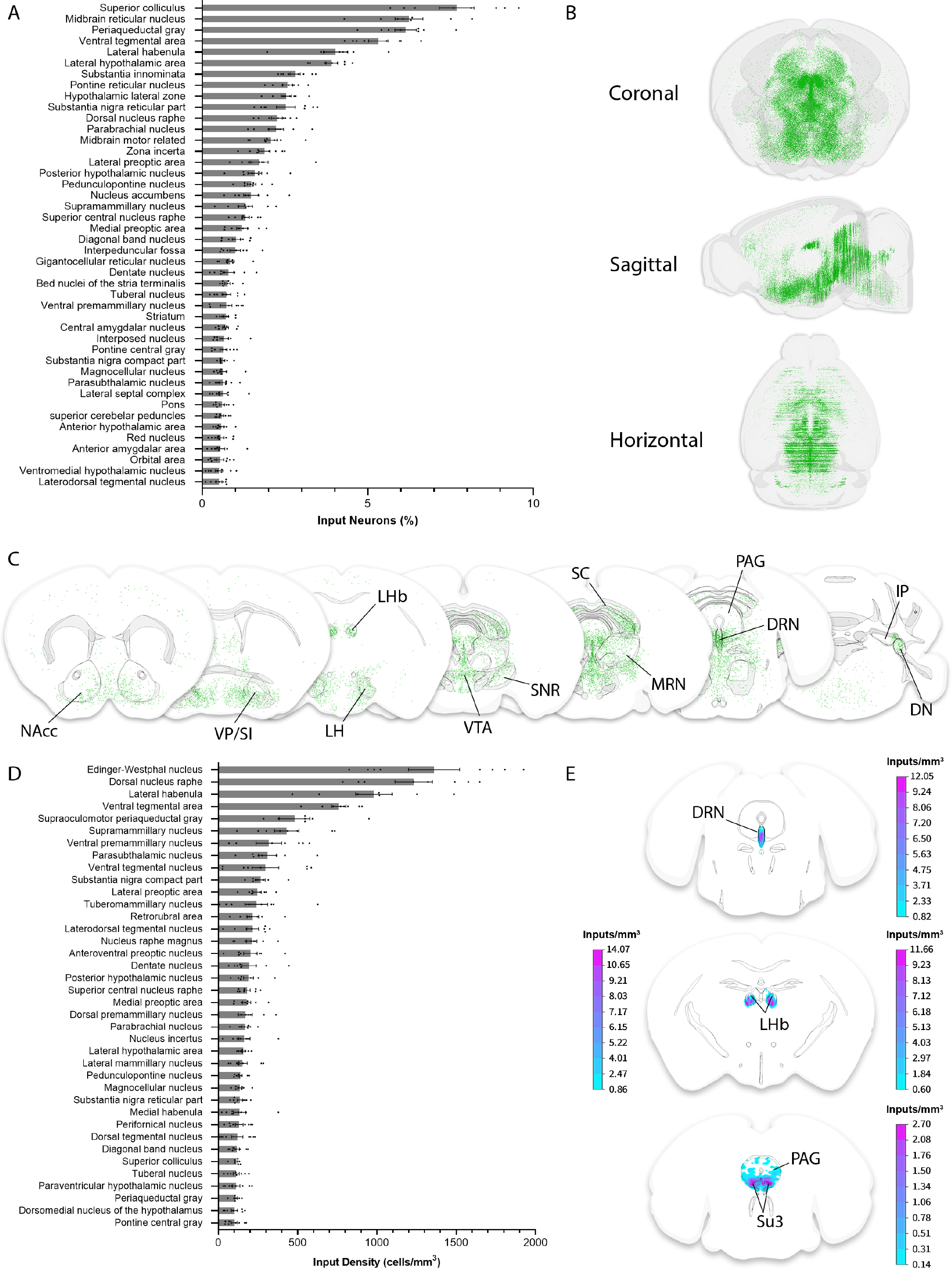
Brain-wide monosynaptic inputs to VTA VGluT2+VGaT+ neurons. **A**. Average proportion of input neurons. All regions accounting for at least 0.5% of inputs are depicted. Dots represent individual animals, *N* = 8. **B**. Coronal, sagittal, and horizontal 3D representations of all input neurons (*N* = 8 animals). **C**. 3D coronal slice representations of input neurons from (left to right) 1.40 to 1.20, 0.20 to 0, −1.60 to −1.80, −3.15 to −3.35, −3.45 to −3.65, −4.30 to −4.50, and −5.8 to −6.0 mm relative to bregma (*N* = 8 animals). **D**. Average density of input neurons. All regions with at least 10 neurons and a density of at least 100 neurons/mm^3^ are depicted. Dots represent individual animals, *N* = 8. **E**. Density heatmaps of input neurons in the DRN (top, −4.20 to −4.40 mm relative to bregma), LHb (middle, −1.60 to −1.80 mm relative to bregma), and PAG (bottom, −3.80 to −4.00 mm relative to bregma). The bilateral LHb has a scale bar for each hemisphere. Abbreviations: DN = dentate nucleus, DRN = dorsal raphe nucleus, IP = interposed nucleus, LH = lateral hypothalamus, LHb = lateral habenula, MRN = midbrain reticular nucleus, NAcc = nucleus accumbens, PAG = periaqueductal gray, SC = superior colliculus, SNR = substantia nigra reticulata, VP/SI = ventral pallidum/substantia innominata, VTA = ventral tegmental nucleus.

Cortical inputs accounted for a substantially smaller proportion of total inputs than subcortical inputs (2.92% of total, 243 ± 28.22 cortical inputs per mouse). Nonetheless, the primary cortical inputs arose from secondary motor cortex (17.51 ± 1.57%), orbital cortex (15.03 ± 3.79%), and insular cortex (14.90 ± 2.06%) (**Figure 4**). Smaller proportions of inputs arose from more sensory-related regions including somatosensory, piriform, gustatory, and visceral cortical regions.

**Figure 4.**
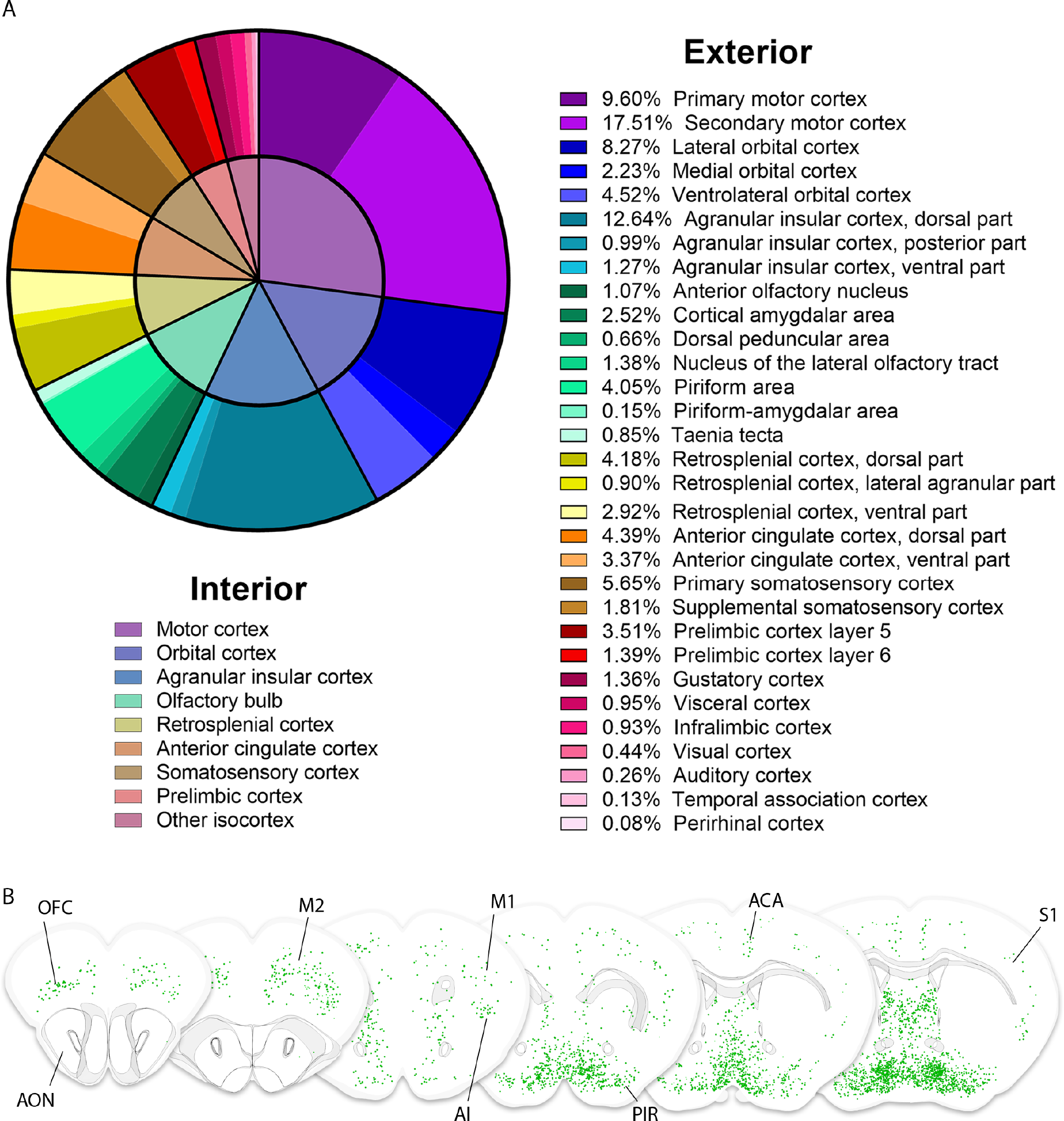
Cortical monosynaptic inputs to VTA VGluT2+VGaT+ neurons. **A**. Average proportion of input neurons in cortical regions. Interior pie chart illustrates broad cortical categories. Exterior pie chart illustrates subregions of cortical categories. **B**. 3D coronal slice representations of input neurons (*N* = 8 animals) from (left to right) 3.00 to 2.80, 2.50 to 2.30, 2.00 to 1.80, 1.50 to 1.30, 1.00 to 0.80, and 0.50 to 0.30 mm relative to bregma. Abbreviations: ACA = anterior cingulate cortex, AI = agranular insular cortex, AON = anterior olfactory nucleus, M1 = primary motor cortex, M2 = secondary motor cortex, OFC = olfactory cortex, PIR = piriform area, S1 = primary somatosensory cortex.

LH has previously been shown to be a primary input to VTA VGluT2+ neurons (Faget et al., 2016; An et al., 2021). Channelrhodopsin2 activation of LH VGluT2+ inputs to VTA increases c-Fos expression in VTA VGluT2+VGaT-neurons but not VGluT2+VGaT+ neurons (Barbano et al., 2020). Nevertheless, given the strong input from LH VGluT2 neurons to VTA VGluT2 neurons (Barbano et al., 2020), we hypothesized that the large number and density of LH neurons that synapse on VTA VGluT2+VGaT+ neurons (**Figure 5A-B**) were glutamatergic. Further, we hypothesized that activation of LH VGluT2 neurons would increase the activity of VTA VGluT2+VGaT+ neurons on a phasic timescale but may not sufficiently increase c-Fos expression. To test this hypothesis, VGluT2::Cre/VGaT::Flp mice were injected in LH with a Cre-dependent vector that expressed the red-shifted opsin ChRmine in LH VGluT2+ neurons and injected in VTA with a Cre- and Flp-dependent vector that expressed GCaMP6m in VTA VGluT2+VGaT+ neurons (**Figure 5C**; **Figure 6A-D**). ChRmine activation of LH VGluT2 neurons robustly increased VTA VGluT2+VGaT+ neuron activity (main effect of stimulation over baseline: *F*(1,3) = 11.657, *p* = 0.042), which was invariant to stimulation frequency (*F*(3,9) = 1.485, *p* = 0.283) (**Figure 5D**). At stimulation onset, we noticed that mice frequently attempted to rear or tread within the enclosed environment used for frequency testing. To more closely examine the behaviors induced by LH VGluT2 neuron stimulation and determine if this behavioral response is related to VTA VGluT2+VGaT+ neuron activity, we stimulated LH VGluT2 neurons while mice were tail-restricted in an unenclosed environment. We chose 20 Hz stimulation because this frequency of activating LH VGluT2 neurons causes defensive behavior that depends on VTA VGluT2+ neurons (Barbano et al., 2020). In the unenclosed environment, stimulation of LH VGluT2 neurons at 20 Hz resulted in increased rearing and paw treading behavior in tail-restricted mice (**Figure 5E-F**; **Movie 1**) as well as increased VTA VGluT2+VGaT+ neuronal activity (**Figure 5G**). However, there was no significant correlation between LH VGluT2 ChRmine-induced reaction durations and VTA VGluT2+VGaT+ neuronal activity (maximum z-score or area under the curve, all *R*^2^ < 0.4, all *p* > 0.07).

**Figure 5.**
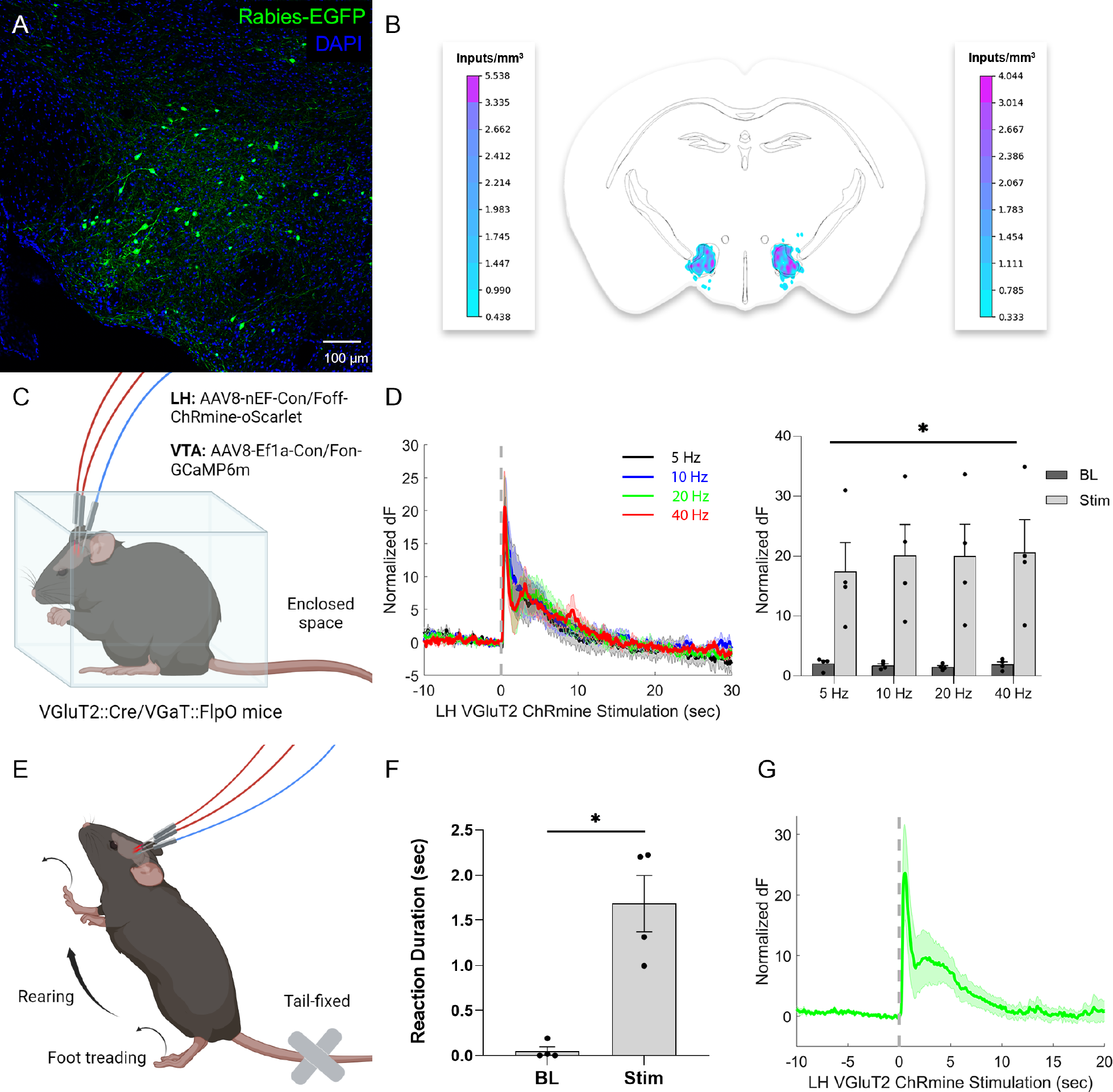
LH VGluT2+ neurons form a functional connection with VTA VGluT2+VGaT+ neurons and cause ambulatory behavior. **A**. Monosynaptic input neurons in LH. **B**. Bilateral density heatmap of input neurons in the LH from −1.60 to −1.80 mm relative to bregma. Left scale bar corresponds to left hemisphere; right scale bar corresponds to right hemisphere. **C-D**. Mice were first optically stimulated in an enclosed space. **C**. Schematic of viral injections and bilateral optical fiber implants for simultaneous optical stimulation of LH VGluT2+ neurons and fiber photometry of VTA VGluT2+VGaT+ neurons. **D**. Normalized averages (left) of GCaMP response to different frequencies of ChRmine laser stimulation. Average z-scores (right) at stimulation were significantly higher than baseline, invariant to stimulation frequency. **E-I**. Mice were then optically stimulated in an unenclosed space while tail-restricted. **E**. Depiction of rearing and foot treading behavior elicited by ChRmine stimulation. **F**. Rearing and foot treading were significantly higher immediately following ChRmine stimulation compared to before stimulation. **G**. Normalized average GCaMP response to 20 Hz ChRmine stimulation over 10 trials (*N* = 4 animals).

**Figure 6.**
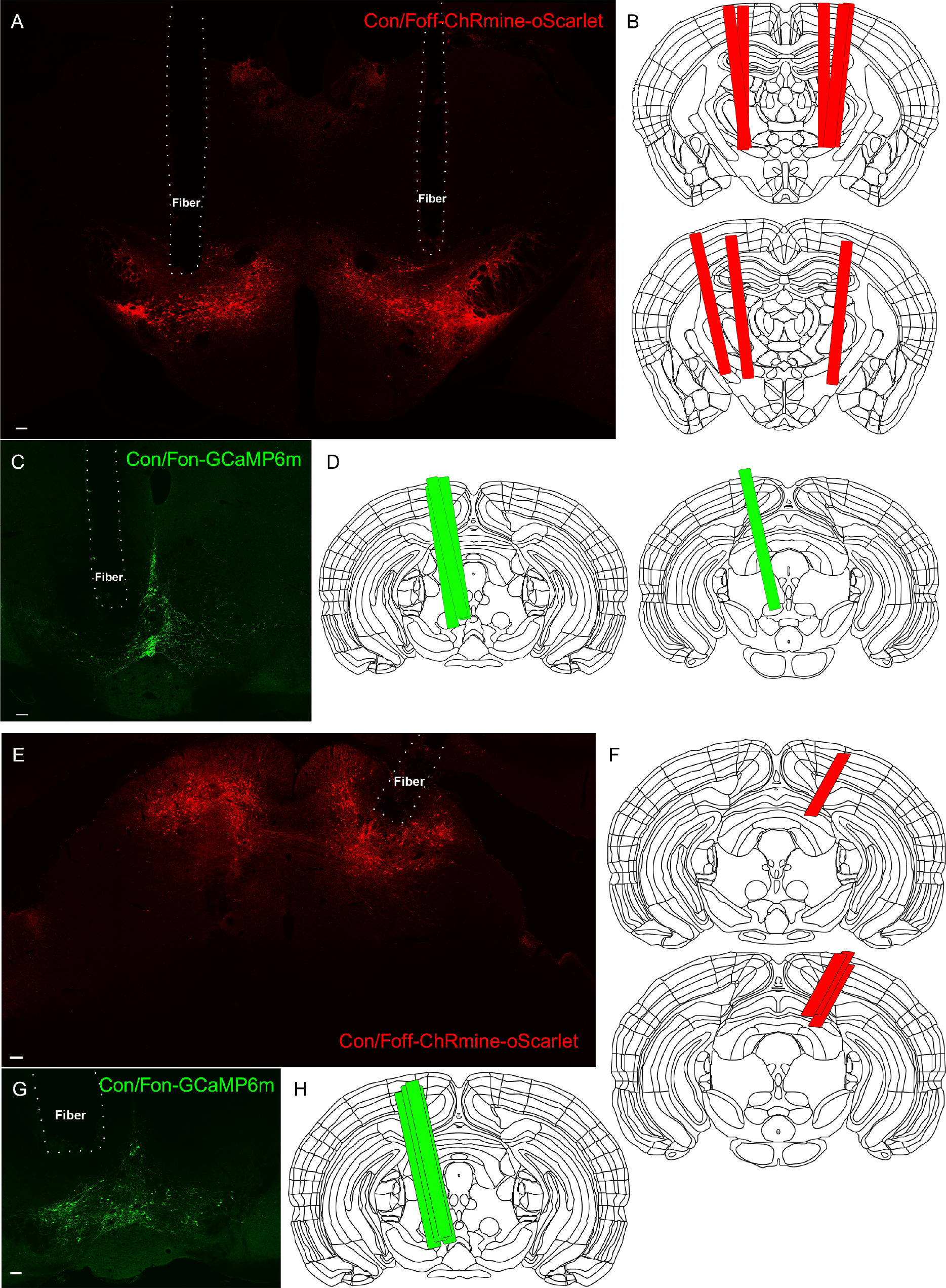
Histological localization of fiber optics targeting LH or SC VGluT2 neurons and VTA VGluT2+VGaT+ neurons. **A**. Example bilateral optical fiber placement and viral expression of AAV8-nEF-Con/Foff-ChRmine-oScarlet. **B**. Anatomical locations of all bilateral optical fiber cannulae, *N* = 4 animals. **C**. Example recording fiber placement and viral expression of AAV8-EF1-Con/Fon-GCaMP6m. **D**. Anatomical locations of all medial recording fiber cannulae, *N* = 4 animals. **E**. Example unilateral optical fiber placement and viral expression of AAV8-Nef-Con/Foff-ChRmine-oScarlet. **F**. Anatomical locations of all unilateral optical fiber cannulae, *N* = 4 animals. **G**. Example recording fiber placement and viral expression of AAV8-EF1-Con/Fon-GCaMP6m. **H**. Anatomical locations of all medial recording fiber cannulae, *N* = 4 animals. Scale bars in A,C = 100 µm.

Another major input to VTA VGluT2+ neurons is from the superior colliculus (An et al., 2021) (SC), which accounted for the largest proportion of inputs to VGluT2+VGaT+ neurons (Figure 2A; **Figure 7A**). SC inputs were densest in the intermediate and ventral layers of SC (**Figure 7B**), suggesting SC inputs were from glutamatergic neurons (Masullo et al., 2019; Liu et al., 2022). To test this, VGluT2::Cre/ VGaT::Flp mice were injected in SC with a Cre-dependent vector that expressed the red-shifted opsin ChRmine in SC VGluT2 neurons and injected in VTA with a Cre- and Flp-dependent vector that expressed GCaMP6m in VTA VGluT2+VGaT+ neurons (**Figure 7C; Figure 6E-H**). In contrast to LH VGluT2 inputs, ChRmine activation of SC VGluT2 neurons resulted in frequency-dependent signaling in VTA VGluT2+VGaT+ neurons, *F*(3,9) = 3.8723, *p* = 0.0497. ChRmine activation of SC VGluT2 neurons significantly increased VTA VGluT2+VGaT+ neuronal activity at 10, 20, and 40 Hz, but not 5 Hz (Sidak adjusted pairwise comparisons 5 Hz: *p* = 0.4181, 10 Hz: *p* = 0.0307, 20 Hz: *p* = 0.0009, 40 Hz: *p* = 0.0034) (**Figure 7D**). We noticed that stimulation of SC VGluT2 neurons often resulted in a rapid head movement and body posture change followed by prolonged freezing behavior within the enclosed testing environment.

**Figure 7.**
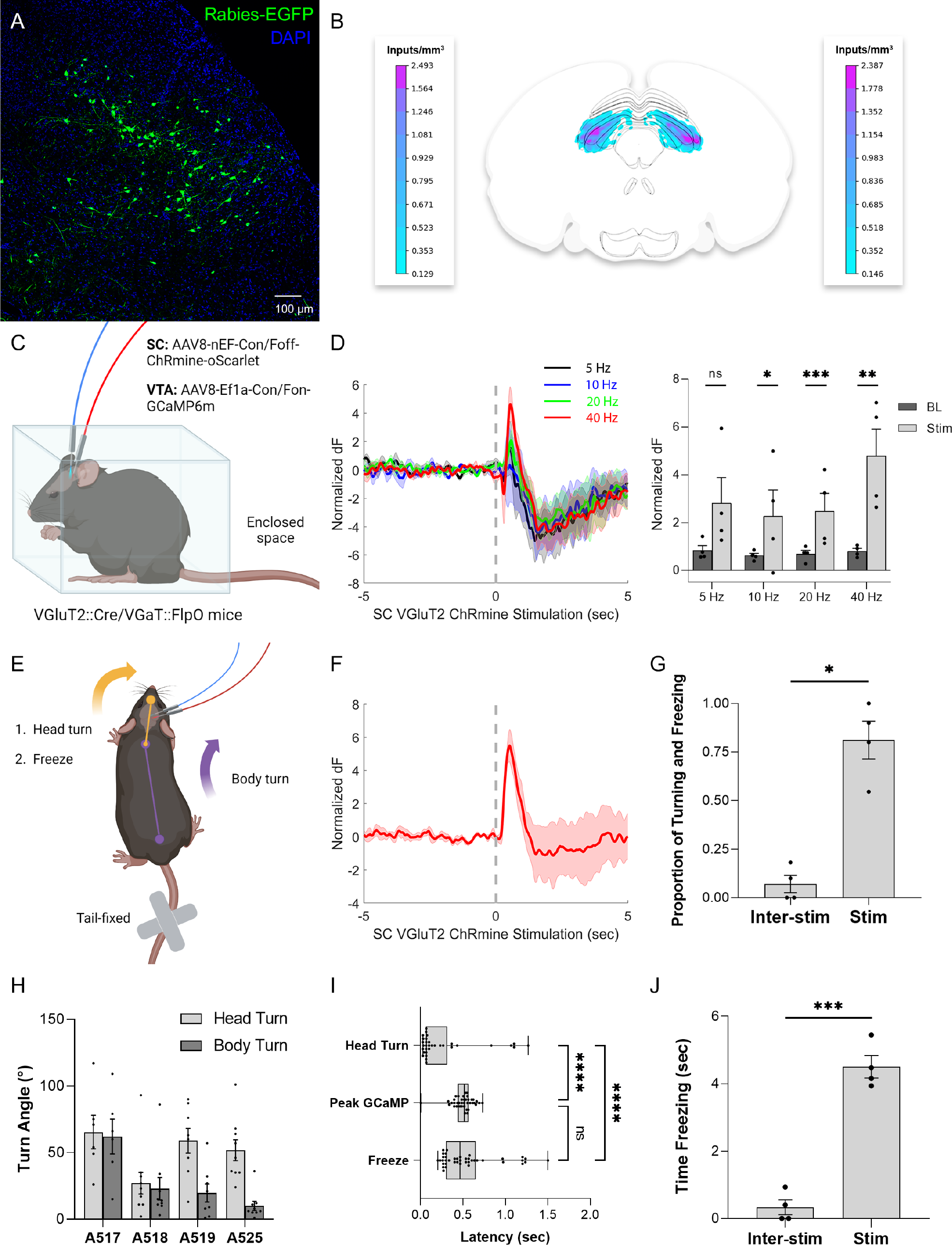
SC VGluT2+ neurons form a functional connection with VTAVGluT2+VGaT+ neurons and reduce ambulatory behavior. **A**. Monosynaptic input neurons in the SC. **B**. Bilateral density heatmap of input neurons in the SC from −3.50 to −3.70 mm relative to bregma. Left scale bar corresponds to left hemisphere; right scale bar corresponds to right hemisphere. **C-D**. Mice were first optically stimulated in an enclosed space. **C**. Schematic of viral injections and unilateral optical fiber implant for simultaneous optical stimulation of SC VGluT2+ neurons and fiber photometry of VTA VGluT2+VGaT+ neurons. **D**. Normalized averages (left) of GCaMP response to different frequencies of ChRmine laser stimulation. Average z-scores (right) depended on the interaction of frequency and stimulation condition. **E-J**. Mice were then optically stimulated in an unenclosed space while tail-restricted. **E**. Depiction of head turning, freezing, and body turning behavior elicited by ChRmine stimulation. **F**. Normalized average GCaMP response to 40 Hz ChRmine stimulation over 10 trials (*N* = 4 animals). **G**. Proportion of stimulation trials and inter-stimulation intervals that resulted in head turning with freezing. Dots represent individual animals. Head turning and freezing behavior was significantly more likely after stimulation compared to during inter-stimulation intervals. **H**. Average angle of head turn and body turn from starting position to freezing position following ChRmine stimulation for each test subject. Dots represent individual stimulation trials. All subjects except A518 had head turn angles normally distributed around their mean turn angle. Subjects A517 and A519 had body turn angles normally distributed around their mean turn angle, while A518 and A525 had body turn angle distributions that departed from normality. **I**. Box-and-whisker plot of head turn, peak GCaMP, and freeze latencies following ChRmine stimulation. Dots represent individual stimulation trials. Head turn latency was significantly shorter than peak GCaMP latency and freeze latency. Peak GCaMP and freeze latency did not differ. **J**. Average time spent freezing immediately following ChRmine stimulation and during inter-stimulation intervals, irrespective of pairing with head turning. The duration of continuous freezing bouts following stimulation was significantly longer than freezing in inter-stimulation intervals.

To more closely examine the behaviors induced by SC VGluT2 stimulation and determine if this behavioral response was related to VTA VGluT2+VGaT+ neuron activity, we stimulated SC VGluT2 neurons at the most reliable frequency that activated VTA VGluT2+VGaT+ neurons (40 Hz) while mice were tail-restricted in an unenclosed environment. In the unenclosed environment, SC VGluT2 neuron stimulation at 40 Hz caused a constellation of behaviors (**Figure 7E**; **Movie 2**) and again activated VTA VGluT2+VGaT+ neuronal activity (**Figure 7F**). ChRmine stimulation resulted in head turning followed immediately by freezing, which was rarely observed during inter-stimulation intervals, *t*(3) = 5.278, *p* = 0.0133 (**Figure 7G**). Following ChRmine stimulation, mice turned their heads an average of 48.45° ± 5.07° from their starting position to their freezing position. All subjects except one (subject A518: Shapiro-Wilk test, *W* = 0.76, *p* = 0.0042) had head turn angles normally distributed around their mean turn angle. Body turns were observed with less vigor and consistency compared to head turns. The mean angle of body turn following SC VGluT2 ChRmine stimulation was 25.78° ± 4.92°. Two of four subjects had body turn angles that departed from a normal distribution around their mean angle (subject A518: Shapiro-Wilk test, W = 0.68, p = 0.0009; subject A525: W = 0.69, p = 0.0012) (Figure 5H). A Kruskal-Wallis test yielded overall differences in the mean latencies to head turn (0.26 ± 0.06 sec after SC VGluT2 stimulation onset), freezing behavior (0.56 ± 0.05 sec), and maximum VTA VGluT2+VGaT+ GCaMP activity (0.50 ± 0.02 sec) (H = 29.30, p < 0.0001). Dunn’s multiple comparison tests found that this main effect was explained by significant differences between head turn latency and freeze latency (p < 0.0001), and between head turn latency and time to peak GCaMP activity (p <0.0001), but not between freeze latency and time to peak GCaMP (p = 1.0) (**Figure 7I**). Irrespective to pairing with head turning, freezing behavior was rarely observed in the inter-stimulation intervals. The duration of the first continuous freezing bout following SC VGluT2 ChRmine stimulation was significantly longer than all freezing in inter-stimulation intervals, *t*(3) = 18.44, *p* = 0.0003 (**Figure 7J**). Subject A518 displayed moderate negative correlations between head turn angle and maximum GCaMP value (*R*^2^ = 0.508, *p* = 0.0207), head turn angle and GCaMP AUC (*R*^2^ = 0.610, *p* = 0.0076), and freeze duration and maximum GCaMP value (*R*^2^ = 0.450, *p* = 0.0339). On the whole, however, there was no significant correlation between SC VGluT2 ChRmine-induced behaviors (latency to head turn, head turn angle, latency to freeze, freeze duration) and VTA VGluT2+VGaT+ neuronal activity (maximum z-score or area under the curve, all other *R*^2^ < 0.6, all other *p* > 0.07). Unlike other VTA neurons, VTA VGluT2+VGaT+ neurons were recently classified by their sensitivity to outcomes but not the learned predictors of those outcomes (Root et al., 2020). VGluT2::Cre/VGaT::Flp GCaMP mice that were previously stimulated in LH or SC VGluT2 neurons were trained to associate an auditory CS+ with footshock over three days, and GCaMP recordings of VTA VGluT2+VGaT+ neurons were conducted on a final day of CS+ footshock pairings (**Figure 8A**). In a separate group of wildtype mice, this training resulted in significantly increased CS+ induced freezing over baseline pre-cue freezing levels, *t*(9) = −2.379, *p* = 0.041 (**Figure 8B**). VTA VGluT2+VGaT+ neuronal activity differed between baseline, cue, and shock epochs with a main effect of time, *F*(2,14) = 7.673, *p* = 0.027. Pairwise comparisons showed that footshock significantly increased VTA VGluT2+VGaT+ neuronal activity over baseline, F(1,7) = 7.684, *p* = 0.028, but there was no difference in activity between baseline and CS+, *F*(1,7) = 0.205, *p* = 0.664 (**Figure 8C-E**). We then assessed whether a relationship existed between LH VGluT2 or SC VGluT2 stimulation-induced VTA VGluT2+VGaT+ neuronal activity and footshock-related activity of VGluT2+VGaT+ neurons. We compared 40 Hz stimulation because both LH VGluT2 and SC VGluT2 stimulation significantly maximally-increased VGluT2+VGaT+ neuronal activity at this frequency. While SC VGluT2 neuron stimulation was not correlated with VGluT2+VGaT+ footshock-induced neuronal activity, *r* = 0.448, *p* = 0.552, LH VGluT2 neuron stimulation was highly correlated with VGluT2+VGaT+ footshock-induced neuronal activity, *r* = 0.983, *p* = 0.017 (**Figure 8F**). Together, these results support that VTA VGluT2+VGaT+ neurons signal outcomes, but not their learned predictors, and suggest that LH VGluT2 neurons play a role in VTA VGluT2+VGaT+ signaling of aversive outcomes.

**Figure 8.**
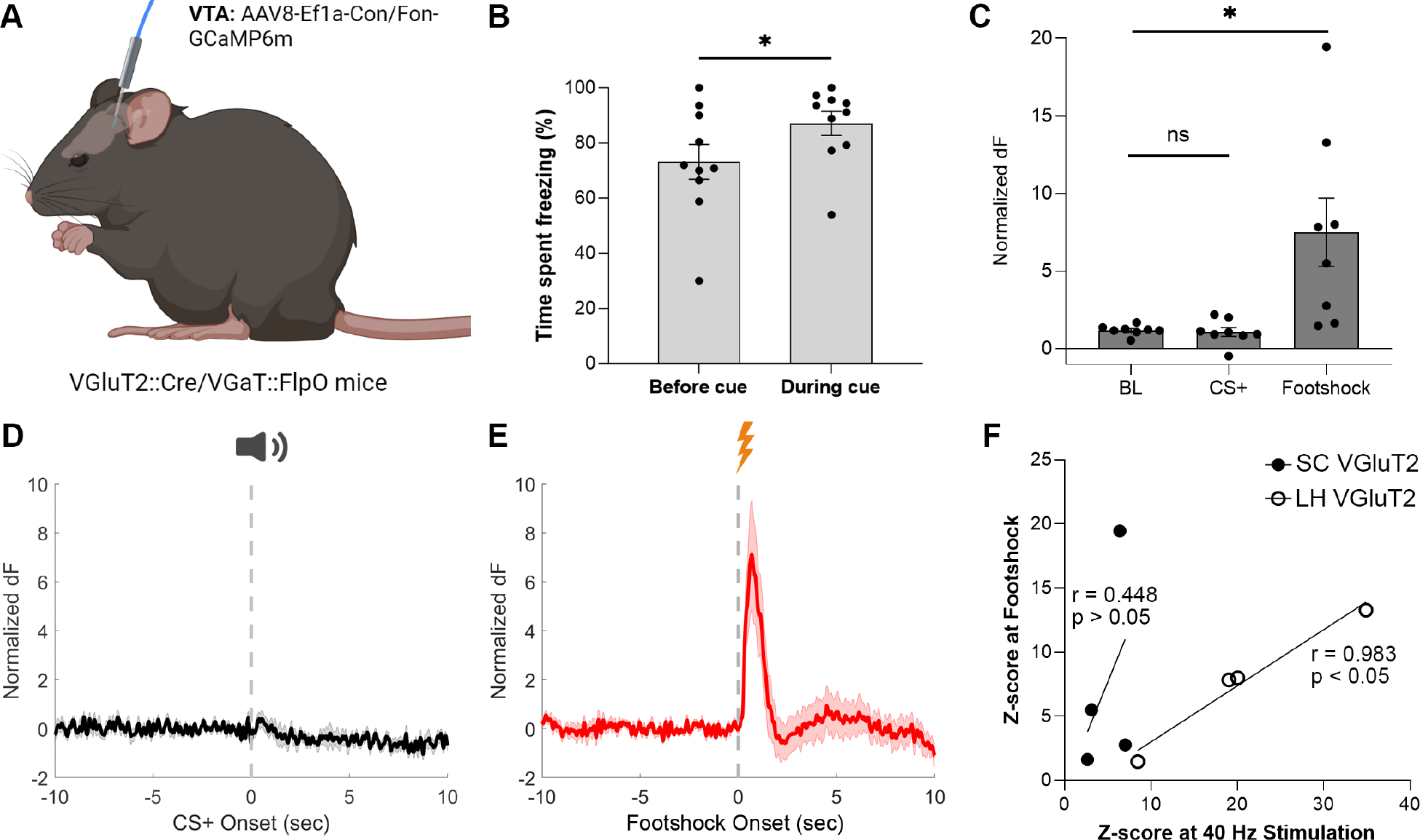
VTA VGluT2+VGaT+ neurons signal footshock outcomes that are correlated with lateral hypothalamus VGluT2+ stimulation-induced VGluT2+VGaT+ activity. **A**. Schematic of viral injection and optical fiber implant for fiber photometry of VTA VGluT2+VGaT+ neurons from mice expressing ChRmine in either lateral hypothalamus (LH) VGluT2 neurons or superior colliculus (SC) VGluT2 neurons. **B**. Wildtype mice show increased freezing behavior in response to a footshock-predicting CS+ compared with prior to CS+ presentation. **C**. Normalized average maximum GCaMP response at baseline, CS+, and footshock. Maximum GCaMP response was significantly higher at footshock compared with baseline and CS+. Dots represent individual animals, *N* = 8. **D**. Normalized average GCaMP response at CS+ onset. **E**. Normalized average GCaMP response at footshock onset. **F**. Correlation between VTA VGluT2+VGaT+ footshock-induced neuronal activity and 40 Hz ChRmine stimulation-induced neuronal activity from SC VGluT2 neurons or LH VGluT2 neurons.

## DISCUSSION

It is well established that subsets of VTA neurons are capable of releasing one or more neurotransmitters (Stuber et al., 2010; Tritsch et al., 2012; Root et al., 2014b; Zhang et al., 2015). Based on the capability of VTA neurons to release one or more neurotransmitters, we hypothesized that the circuits and behavioral functions of VTA neurons depend on multiple genetic characteristics of neurotransmitter release. In support, VTA neurons that release glutamate without GABA (VGluT2+VGaT-), release GABA without glutamate (VGluT2-VGaT+), release both glutamate and GABA (VGluT2+VGaT+), or release dopamine (TH+) have distinct and partially overlapping axonal targets as well as unique neuronal activation profiles in response to rewarding or aversive stimuli and their learned predictors (Root et al., 2020). Here, we focused on the population of VTA neurons that release both glutamate and GABA (VGluT2+VGaT+ neurons) and are activated by rewarding or aversive outcomes but not learned predictors of those outcomes (Root et al., 2020). We aimed to identify the circuit inputs that regulate VTA VGluT2+VGaT+ neuronal signaling and their contribution to behavior, which are currently unknown.

Monosynaptic rabies viral tracing with helper viruses dependent on Cre and Flp recombinases was used to identify brainwide inputs to VTA VGluT2+VGaT+ neurons. To determine the intersectional specificity of the rabies helper viruses, we injected VGluT2::Cre mice and VGaT::Flp mice with the same Cre and Flp-dependent AAVs encoding TVA-mCherry and oG, followed by rabies-EGFP three weeks later. While we did not observe mCherry-labeled neurons or retrogradely labeled EGFP neurons throughout the brain, we observed EGFP-labeled neurons locally within the injection site. The number of locally labeled EGFP neurons was small, approximately 2.55% of starter neurons in VGluT2::Cre/VGaT::Flp mice. Local EGFP labeling may have been due to leaky recombinase-independent expression of TVA and the phenomenon termed “invisible TVA” (Hafner et al., 2019). In the original description of intersectional rabies virus tracing, the authors report similar levels of leaky TVA expression in mice expressing only one of the two required recombinases, with an average of 10.0 and 4.0 rabies-EGFP cells in Cre-only mice and Flp-only mice, respectively. Furthermore, starter cells at the injection site may be “invisible” by expressing enough TVA-mCherry to allow the rabies-EGFP virus to transduce the cell, but not enough to express detectable levels of mCherry fluorophore. These cells would therefore appear to express only EGFP, and would be presumed input neurons rather than starter neurons. The lack of rabies-EGFP cells outside the injection site supports leaky and invisible TVA but not leaky oG, since oG would allow rabies virus to spread transsynaptically from the starter cells, which was not observed in our single recombinase controls. Taken together, these results indicate minor off-target rabies-EGFP labeling of neurons likely expressing only Cre or only Flp in the injection site due to leaky, invisible TVA. Accordingly, the number of local VTA input neurons reported here may represent an overestimation of 1.51%. However, this off-target labeling at the injection site did not spread transsynaptically, due to the intersectional specificity of oG. This supports the assumption that retrogradely labeled input neurons counted outside the VTA correspond to on-target VGluT2+VGaT+ starter neurons.

Using monosynaptic rabies viral tracing of inputs to VGluT2+VGaT+ neurons, we found that most synaptic inputs arose from subcortical structures implicated in motor actions, outcome valuation, and threat responding, such as superior colliculus, lateral hypothalamus, lateral habenula, periaqueductal gray, dorsal raphe, substantia innominata/ventral pallidum, locally within the VTA, and parabrachial nucleus (Matsumoto and Hikosaka, 2009; Bromberg-Martin et al., 2010; Hikosaka, 2010; Thevarajah et al., 2010; Benarroch, 2012; Cohen et al., 2012; Lecca et al., 2014; Root et al., 2015; Bonnavion et al., 2016; Stuber and Wise, 2016; Basso and May, 2017; Zahm and Root, 2017; Palmiter, 2018; Silva and McNaughton, 2019; Root et al., 2020; Hoang and Sharpe, 2021). These circuits likely contribute to the high sensitivity of VTA VGluT2+VGaT+ neurons toward stressful stimuli that result in social and exploratory deficits in mice (McGovern et al., 2022). On the whole, brain-wide inputs to VTA VGluT2+VGaT+ neurons were similar to those found examining all VTA VGluT2+ neuron inputs (Faget et al., 2016; An et al., 2021). Highly dense inputs arose from lateral habenula, dorsal raphe, locally within the VTA, and subdivisions of the periaqueductal gray. The highest density of inputs was defined by the Allen Atlas as the Edinger-Westphal nucleus. However, this labeled region best corresponds to subdivisions of the VTA involving the linear portions of the rostral linear nucleus and caudal linear nucleus that contain large numbers of VGluT2-expressing neurons (Root et al., 2018b). Further, because Edinger-Westphal nucleus neurons release peptides, but not neurotransmitters such as glutamate or GABA (Priest et al., 2023), it is unlikely that rabies transsynaptic tracing would label Edinger-Westphal nucleus neurons that use paracrine signaling. Thus, we consider the dense inputs within the labeled Edinger-Westphal nucleus as local VTA inputs, which were large in number and density.

Cortically, there were significantly fewer inputs to VTA VGluT2+VGaT+ neurons than subcortical inputs, which is consistent with inputs to other VTA cell-types (Watabe-Uchida et al., 2012; Faget et al., 2016; An et al., 2021). More than half of cortical inputs were from primary and secondary motor cortex, orbital cortex, and agranular insular cortex, which are implicated in action selection, outcome valuation, decision making, and fear memory (Schoenbaum et al., 2011; Sul et al., 2011; Shi et al., 2020; Knudsen and Wallis, 2022). Integration of M2 inputs, the largest cortical input region onto VTA VGluT2+VGaT+ neurons, could represent a proprioceptive feedback circuit to ensure correct responding to relevant outcomes.

Based on our prior examinations of glutamate or GABA inputs to VTA VGluT2 neurons (McGovern et al., 2021) and the location of lateral hypothalamic and superior colliculus inputs to VTA VGluT2+VGaT+ neurons, we hypothesized these pathways were glutamatergic. Optogenetic stimulation of the LH VGluT2 neuron → VTA pathway results in real-time place avoidance and reduction of accumbal dopamine (Nieh et al., 2016). Further, fleeing behavior that is induced by a visual threat depends on a LH VGluT2 neuron → VTA VGluT2 neuron pathway (Barbano et al., 2020). We found that LH VGluT2 neurons robustly increased VTA VGluT2+VGaT+ neuronal activity across frequency parameters. LH VGluT2 stimulation also resulted in rearing and paw treading movements that resembled an attempted flee or escape behavior. However, the activation of VTA VGluT2+VGaT+ neurons by LH VGluT2 neurons was not correlated with the behavioral reactions induced by LH VGluT2 stimulation. The escape-like behaviors following LH VGluT2 neuron activation may arise from LH VGluT2 innervation of VTA VGluT2+VGaT-neurons (Barbano et al., 2020) or other regions such as the lateral habenula (Lecca et al., 2017), of which the LHb was among the densest inputs to VTA VGluT2+VGaT+ neurons. We previously found that VTA VGluT2+VGaT+ neurons are highly sensitive to repeated, uncontrollable tailshocks that induce social avoidance, exaggerated fear, and reduced exploratory behavior (McGovern et al., 2022). Further, lateral habenula neurons receiving synapses from VTA VGluT2+ neurons, which are mostly VGluT2+VGaT+ neurons(Root et al., 2014b), are also highly activated by repeated uncontrollable tailshock stress (McGovern et al., 2022). These results suggest that interactions between LH VGluT2 neurons, lateral habenula neurons, VTA VGluT2+VGaT-, and VTA VGluT2+VGaT+ neurons may be important for immediate threat-related actions as well as long-term repercussions of prior threat experience.

Optogenetic stimulation of SC VGluT2 neurons increased VTA VGluT2+VGaT+ neuron activity at comparatively higher frequencies than LH VGluT2 neurons. Previous research has shown that optogenetic stimulation of the SC VGluT2 → VTA pathway activates VTA dopamine neurons as well as VTA GABA neurons within the medial VTA where VGluT2+VGaT+ neurons are located, but not laterally where VGluT2-VGaT+ neurons predominate (Root et al., 2018b; Root et al., 2020; Solie et al., 2022). We found that SC VGluT2 stimulation resulted in head turning followed by freezing, which resembled orientation toward and avoidance of detection by a threat (Dean et al., 1989; Bradley, 2009). In support of the role of the SC VGluT2 → VTA pathway in signaling orientation toward threatening stimuli, the densest SC inputs were consistent with the location of glutamatergic Pitx2 SC neurons which drive head orientation (Masullo et al., 2019). However, head turns occurred before the peak activation of VTA VGluT2+VGaT+ neurons, and the activity of VGluT2+VGaT+ neurons did not correlate with movement parameters of the head turn, indicating VTA VGluT2+VGaT+ neurons are likely unnecessary for generating this head orientation response. Likewise, although the peak activation of VTA VGluT2+VGaT+ neurons was concurrent with freezing behavior, there was no correlation between VGluT2+VGaT+ activity and freezing behavior. Together with the results of LH VGluT2 neuronal stimulation, we found no evidence that the outcome signaling of VTA VGluT2+VGaT+ neurons is directly related to movement kinematics during outcomes. Given that both LH VGluT2 and SC VGluT2 neurons activated VTA VGluT2+VGaT+ neurons, and LH VGluT2 neurons caused ambulatory escape-like behavior while SC VGluT2 neurons caused halted ambulatory behavior (freezing), a common feature of VTA VGluT2+VGaT+ neuronal activation between pathways is a behavioral response consistent with the interpretation of a threat.

In conclusion, these results provide novel insights into the cell-type specific influences on VTA VGluT2+VGaT+ neuronal activity. We found that VTA VGluT2+VGaT+ neurons received large numbers of monosynaptic inputs from SC, LH, midbrain reticular nucleus, and periaqueductal gray, whereas the densest inputs were from dorsal raphe nucleus, LHb, and VTA. We also found that stimulation of LH VGluT2 and SC VGluT2 neurons each activated VTA VGluT2+VGaT+ neurons *in vivo*. LH stimulation more reliably resulted in activation of VGluT2+VGaT+ neurons across frequencies compared with SC stimulation, which activated VGluT2+VGaT+ neurons at high frequency. Our initial examination of these pathways revealed that LH VGluT2 and SC VGluT2 stimulation induced opposite ambulatory behavioral responses, and while some threat-response behaviors were concurrent with VGluT2+VGaT+ neuronal activity, no stimulation-induced behavioral kinematics correlated with VTA VGluT2+VGaT+ neuronal activity. Instead, we found that LH VGluT2 stimulation-induced activation of VTA VGluT2+VGaT+ neurons was highly correlated with footshock-induced activation of VTA VGluT2+VGaT+ neuronal activity, while SC VGluT2 stimulation did not correlate with footshock-induced activity. We interpret these data such that we did not find evidence to support the role of VTA VGluT2+VGaT+ neurons in generating immediate threat-related kinematics. The immediate threat-related kinematics may result from other projections of LH VGluT2 or SC VGluT2 neurons whereas a generalized threat signal is integrated onto VTA VGluT2+VGaT+ neurons regardless of the motor behaviors that underlie them. These inputs may specialize, because footshock-related activation of VTA VGluT2+VGaT+ was highly correlated with LH glutamatergic input while not correlated with SC glutamatergic input. Nevertheless, the role of these neurons as integrators of signals related to aversive outcomes is supported by their primary output to LHb (Root et al., 2014b; Root et al., 2018b; Root et al., 2020). The high degree of reciprocal connectivity of VTA VGluT2+VGaT+ neurons, LH VGluT2 neurons, and LHb neurons could suggest extensive processing, amplification, or filtering of aversive signaling. However, the high sensitivity of VTA VGluT2+VGaT+ neurons to rewarding outcomes (Root et al., 2020) suggests a wider role in outcome detection and salience processing that likely arises from regions outside of the LH and SC.

## ACKNOWLEDGEMENTS

This research was supported by the Webb-Waring Biomedical Research Award from the Boettcher Foundation (DHR), R01 DA047443 (DHR), F31 MH125569 (DJM), a 2020 NARSAD Young Investigator grant from the Brain and Behavior Research Foundation (DHR), The Curci Foundation (EDP), and The University of Colorado Boulder. The imaging work was performed at the BioFrontiers Institute Advanced Light Microscopy Core (RRID: SCR_018302). Laser scanning confocal microscopy was performed on a Nikon A1R microscope supported by NIST-CU Cooperative Agreement award number 70NANB15H226. The PerkinElmer Opera Phenix is supported by NIH grant 1S10OD025072. The funders had no role in study design, data collection and analysis, decision to publish, or preparation of the manuscript. Prism, MATLAB, Brainrender, and BioRender. com were used to make figures and schematics. We thank Nicholas Steinmetz and Jun Takatoh for their help in density analyses.

The authors report no biomedical financial interests or potential conflicts of interest.

**Movie 1. LH VGluT2+ neuronal activation causes escape-like behavior**. VGluT2::Cre/VGaT::Flp mice with ChRmine expression and bilateral optic fiber cannulae in LH. While tail-restricted in an unenclosed environment, a one-second train of 20 Hz ChRmine stimulation induced foot treading (view from below) and rearing (view from the side) onto the hind legs. Responses began shortly after laser onset and continued for an average of 1.74 ± 0.14 s.

**Movie 2. SC VGluT2+ neuronal activation causes behavior resembling threat detection and avoidance**. VGluT2::Cre/ VGaT::Flp mouse with ChRmine expression and an optic fiber cannula in SC, view from above. While tail-restricted in an unenclosed environment, a one-second train of 40 Hz ChRmine stimulation induced head orientation followed by a prolonged freezing bout. Head turn began immediately after laser onset. Mice remained frozen for an average of 4.52 ± 1.68 s.

